# Deep sampling and pooled amplicon sequencing reveals hidden genic variation in heterogeneous rye accessions

**DOI:** 10.1101/2020.02.20.958181

**Authors:** Anna Hawliczek, Leszek Bolibok, Katarzyna Tofil, Ewa Borzęcka, Joanna Jankowicz-Cieślak, Piotr Gawroński, Adam Kral, Bradley J Till, Hanna Bolibok-Brągoszewska

## Abstract

Loss of genetic variation negatively impacts breeding efforts and food security. Genebanks house over 7 million accessions representing vast allelic diversity that is a resource for sustainable breeding. Discovery of DNA variations is an important step in the efficient use of these resources. While technologies have improved and costs dropped, it remains impractical to consider resequencing millions of accessions. Candidate genes are known for most agronomic traits, providing a list of high priority targets. Heterogeneity in seed stocks means that multiple samples from an accession need to be evaluated to recover available alleles. To address this we developed a pooled amplicon sequencing approach and applied it to the out-crossing cereal rye *(Secale cereale).* Ninety-five rye accessions of different improvement status and worldwide origin, each represented by a pooled sample comprising DNA of 96 individual plants, were evaluated for sequence variation in six target genes involved in seed quality, biotic and abiotic stress resistance. Seventy-four predicted deleterious variants were identified using multiple algorithms. Rare variants were recovered including those found only in a low percentage of seed. A large extent of within-population heterogeneity was revealed, providing an important point for consideration during rye germplasm conservation and utilization efforts. We conclude that this approach provides a rapid and flexible method for evaluating stock heterogeneity, probing allele diversity, and recovering previously hidden variation.

## INTRODUCTION

Plants can be made more resilient, yields stabilized, and nutritional components enhanced through selection and combination of gene variants that control these traits. Crop improvement is therefore dependent on the existence of genetic variability for the trait in question. For the past 10,000 years humans have been selecting and combining genetic variants to improve crops. However, most of the history of crop development was carried out without a knowledge of genetics or DNA, and thus modern cultivars have a relatively narrow genetic base, resulting from bottleneck-like effects of domestication and breeding practices (Purugganan and Fuller, 2009; Mondal *et al*., 2016; Joukhadar *et al*., 2017). Therefore, the allelic variability existing within contemporary cultivars or breeding programs may be insufficient for successful identification of gene variants for satisfactory productivity and resilience of the crop.

Useful alleles conferring important traits that have been lost in modern cultivars may still exist in nature. Plant genetic resources (PGR), such as landraces and wild relatives of crop plants, possess a much higher genetic diversity. While not high yielding and having often undesirable agronomic characteristics, they were shown to contain gene variants that can improve performance of successful modern cultivars (Hoisington *et al*., 2002; McCouch, 2013; Gur and Zamir, 2004; Gamuyao *et al*., 2012).

Luckily, the value of PGR as a reservoir of gene variants was recognised over a hundred years ago (McCouch, 2004) and nowadays there are over 1,700 *ex situ* germplasm collections worldwide, maintaining about 7.4 million accessions. Approximately 62% of these accessions are landraces and wild species (FAO, 2010). Unfortunately, in most cases little is known about the extent and structure of genetic diversity within a given collection. The available data is often limited to passport information, and some phenotypic measurements or DNA marker-based genetic diversity assessment for a subset of accessions. Such information is not sufficient to make an informed choice of PGR for inclusion into a breeding program. Therefore the utilisation of primitive, exotic germplasm in crop improvement is limited (FAO, 2010; McCouch, 2013; Keilwagen *et al*., 2014).

To fully profit from the allelic variation of PGR, methods for efficient and reliable screening of hundreds of accessions to discover useful gene variants are needed. Rapid development of next generation sequencing (NGS) technologies resulted in the establishment of various approaches, which can be used for high-throughput assessment of genic variation, such as whole genome resequencing (WGS) (Li *et al*., 2013; Mehra *et al*., 2015; Zhou *et al*., 2015) and exome capture (Hodges *et al*., 2007; Hussain *et al*., 2018). Unfortunately, these approaches are not yet applied in many species owing to factors including genome size, polyploidy, and associated costs of sequencing and capture probe development. While a future can be envisioned where comprehensive genomic data is available for every accession of every important crop, the current state of technology and funding means that material is prioritized, and compromises made. Insofar as evaluation of WGS data provides information useful for understanding population genetics and evolution, it is expected that only a small fraction of base pairs of a genome are controlling key agronomic traits (Wendel *et al*., 2016) Targeting candidate genes and their regulatory elements provides a tremendous reduction in data collected. Indeed, many studies have revealed quantitative trait loci and associated candidate genes that can be used to identify orthologous sequences in other plants (Nguyen *et al*., 2019). An alternative to whole genome or exome capture sequencing is amplicon sequencing. In this approach, selected genomic regions are first amplified by PCR and then subjected to massively parallel sequencing. Compared to WGS or exome capture, amplicon approaches allow acquisition of a much higher coverage of the selected target bases pairs at a lower sequencing cost. This is because the total yield of the sequencing reaction, in terms of raw bases, is distributed to fewer unique bases of each sample in the pool (e.g. (Sims *et al*., 2014)). One application of amplicon sequencing is the simultaneous genotyping of hundreds of unique samples independently by employing strategies to barcode, or index, each sample uniquely (Campbell *et al*., 2015). In addition to this approach, the high sensitivity of current sequencing technologies enables “ultra deep” methods whereby nucleotide variants can be identified in samples containing pools of mixed genotypes. One example is the detection of rare somatic mutations in human samples (Dou *et al*., 2018). Another example is the use of amplicon sequencing to measure intrahost virus diversity. Researchers showed that a rare Zika virus variant could be detected if present at > 3% in a mixed sample when sequencing coverage was at least 400x (Grubaugh *et al*., 2019). In plants, experiments can be designed to discover rare nucleotide variants present at very low frequencies by screening large populations where genomic DNA has been pooled prior to PCR amplification and sequencing. Screening throughputs are increased and assay costs are reduced, making screening thousands of samples practical. This has been used for recovery of induced point mutations in TILLING by Sequencing assays (Tsai *et al*., 2011). Here, genomic DNAs from different lines harboring induced mutations are pooled, subjected to target-specific PCR and the PCR products are then pooled and sequenced. The method has been used to recover rare mutations in genomic DNA samples pooled from 64 to 256 fold. These studies suggest that variant calling accuracy is improved when using multiple variant calling algorithms(Tsai *et al*., 2011; Pan *et al*., 2015; Gupta *et al*., 2017; Tramontano *et al*., 2019). The approach has been adapted for recovery of natural variation in *Populus nigra, Manihot esculenta* Crantz (cassava), and *Oryza sativa* L., whereby DNAs from different accessions were pooled together prior to PCR and variant discovery. In *P. nigra*, PCR products were prepared from pooled genomic DNA from 64 accessions to identify variants in lignin biosynthesis genes in 768 accessions (Marroni *et al*., 2011) In cassava, DNA from up to 281 accessions were pooled prior to sequencing for variants in starch biosynthesis pathway-related genes and herbicide tolerance genes in 1667 accessions (Duitama *et al*., 2017). In rice, pooling of DNAs prepared from 233 breeding lines was followed by sequencing for variants in starch synthesis genes (Kharabian-Masouleh *et al*., 2011). Pooling of multiple samples from the same species has also been used in studies where WGS has been applied. There are many variations to this methodology that has been termed Pool-seq (Schlötterer *et al*., 2014) This includes cases where, contrary to TILLING assays, multiple individuals with similar genotypes are pooled together to estimate population allele frequencies. In such applications, sequencing coverages can be reduced to save costs, but are insufficient to find rare alleles in one or few individuals in the pool. Sequencing intra-species pools has also been described such as in metagenomics studies (Pereira *et al*., 2018).

Rye (Se*cale cereale L*.) is an outcrossing cereal, popular in Europe and North America, and an important source of variation for wheat breeding due to its high tolerance to biotic and abiotic stresses (Crespo-Herrera LA et al., 2017). Genetically rye is a diploid (n=7), with a large (ca. 8 Gbp) and complex genome (Bartos *et al*., 2008; Rabanus-Wallace *et al*., 2019). There are over 21 thousand rye accessions in genebanks worldwide, approximately 35% of them are landraces and wild species (FAO, 2010). Several studies on genome-wide diversity in rye were published to date (Bolibok-Bragoszewska *et al.*, 2014; Targońska *et al.*, 2016; Maraci *et al.*, 2018; Sidhu *et al.*, 2019). It was shown that accessions from genebanks are genetically distinct from modern varieties, which highlighted the potential of PGR in extending the variability in current rye breeding programs (Bolibok-Bragoszewska *et al.*, 2014; Targońska *et al.*, 2016). To date neither NGS-based targeted amplicon sequencing, nor any other method of gene variant discovery was applied to rye genetic resources.

Although the number of well characterized rye genes is very limited (Gawroński *et al*., 2016), there are important candidate genes to consider. *MATE1* is a gene involved in aluminum (Al) tolerance of rye. Al-toxicity is one of the main constraints to agricultural production on acidic soils, which constitute ca. 50% of the arable land on Earth (Maron *et al*., 2013). Rye is one of the most Al-tolerant cereals, with the degree of tolerance depending on the allelic variant of *MATE1* (Santos *et al*., 2018). TLPs are a family of pathogenesis-related (PR) proteins, involved in fungal pathogen response in many plant species (Zhang *et al*., 2018). FBA is one of the key metabolic enzymes involved in CO2 fixation and sucrose metabolism. *FBA* genes were found to have an important role in regulation of growth and development, and responses to biotic and abiotic stresses, such as chilling, drought and heat (Lv *et al*., 2017; Cai *et al*., 2018). *GSP*-1 genes, belonging to the prolamin superfamily of seed storage proteins, encode precursor proteins, which after post-translational processing give rise to arabinogalactan peptide AGP and the grain softness protein GSP-1 (Wilkinson *et al*., 2017) Secaloindolines, products of genes *Sina* (not analyzed in this study) and *Sinb*, are main components of friabilin - a starch-associated protein fraction of cereal grains (Simeone and Lafiandra, 2005). The wheat orthologues of *Sina* and *Sinb*, called *Pina* and *Pinb*, are key determinants of grain texture, an important breeding trait directly influencing the end-use (Liu *et al*., 2017). *PBF* is an endosperm specific transcription factor involved in the regulation of protein and starch synthesis (Zhang *et al*., 2016). It binds to the prolamine-box motif occurring in promoter regions of multiple cereal seed storage proteins. In barley, SNPs located in *PBF* were associated with crude protein and starch content (Haseneyer *et al*., 2010), while in wheat, mutating the homeologous *PBF*s using TILLING resulted in a markedly decreased gluten content and high content of lysine (Moehs *et al*., 2019).

Exploration of genic variation in outcrossing, generatively propagated crops, such as rye, maize, sugar beet, broccoli, or carrot, is a particularly demanding task. Natural, random-mating populations of such species are heterozygous and heterogeneous, with multiple alleles of a locus being present (Souza Jr., 2012). Such population structure has important implications for the design of NGS-allele mining experiments. Firstly, due to high levels of heterozygosity, a higher sequencing coverage is needed even when sequencing non-pooled samples to ensure reliable nucleotide variant calling. Secondly, due to the heterogeneity of accessions, a large enough number of individuals of a given accession needs to be included in the screen to obtain a faithful representation of within-accession variability and to successfully recover rare variants. Many potentially useful and interesting alleles may go undiscovered with current experimental designs.

To address this, a low-cost, high-throughput, and reliable amplicon sequencing approach suitable for assessment of genic variation in heterozygous and heterogeneous rye accessions was developed. Rather than pool DNA from different accessions, ultra deep amplicon sequencing was used to evaluate intra-accession heterogeneity while also providing information on novel genetic variation. DNA pools were created that contain ninety-six plants per accession. These were subjected to pooled amplicon sequencing in six target genes implicated in biotic and abiotic stress resistance and seed quality: *MATE1*, *TLP*, *FBA*, *PBF*, *Sinb*, and *GSP-1*. Three variant calling algorithms (GATK HaplotypeCaller (Poplin *et al*., 2017), SNVer (Wei *et al*., 2011) and CRISP (Bansal, 2010)) were used to identify putative variants at frequencies as low as one heterozygous event per 96 plants assayed in each pool. A subset of variants was independently validated and the functional effect of each variant was evaluated *in silico*. Common and rare variants were recovered, including variants predicted to affect protein function that are present in only a small fraction of seed representing an accession. This data provides preliminary knowledge on the levels of variant allele frequencies in accessions representing different germplasm groups: wild species, landraces, historical and modern cultivars.

## RESULTS

### DNA sequencing, mapping and coverage

Pooled amplicon sequencing using Illumina sequencing by synthesis 2×300 paired end reads on 95 accessions and six genes produced a mean coverage of 13948× and mean mapping quality of 58.65. Mean coverage per accession pool varied approximately 10 fold, between 2924× and 30275×. Analysis of sequencing coverage at each nucleotide revealed that 94.2% of the experiment produced 20 or more reads to support a rare variant present at 5% in the DNA pool (Supplemental Table 1).

### Evaluation of variant calling algorithms and predicted effects of nucleotide changes

Variant calling was first performed on each pool using HaplotypeCaller in GATK (v.4.0) with ploidy set to 192 in order to recover rare alleles. This resulted in 4,115 called variants, of which 3,682 were single nucleotide polymorphisms, 192 insertions, and 241 deletions. Evaluation of the Variant Call Format (VCF) file, allowed calculation of the frequency of a specific allele within the DNA pool created from the 96 seeds that were sampled to represent an accession. This is referred to as VAF (Variant Allele Frequency), to distinguish the measurement from AF (Allele Frequency) - the frequency of the allele within the set of 95 accessions analyzed in the present study. Data was plotted to evaluate the distribution of the mean VAF for each variant and the number of accessions harboring each discovered allele (Figure 1). Private variants occurring in only one accession were identified at both low and high VAFs (Figure 1). The percentage was highest, however, at the lowest VAFs - seventy-five percent of private alleles have a VAF of 0.026 (represented at 2.6% in the accession pool) or lower (Supplemental Figure 1).

**Figure 1.**
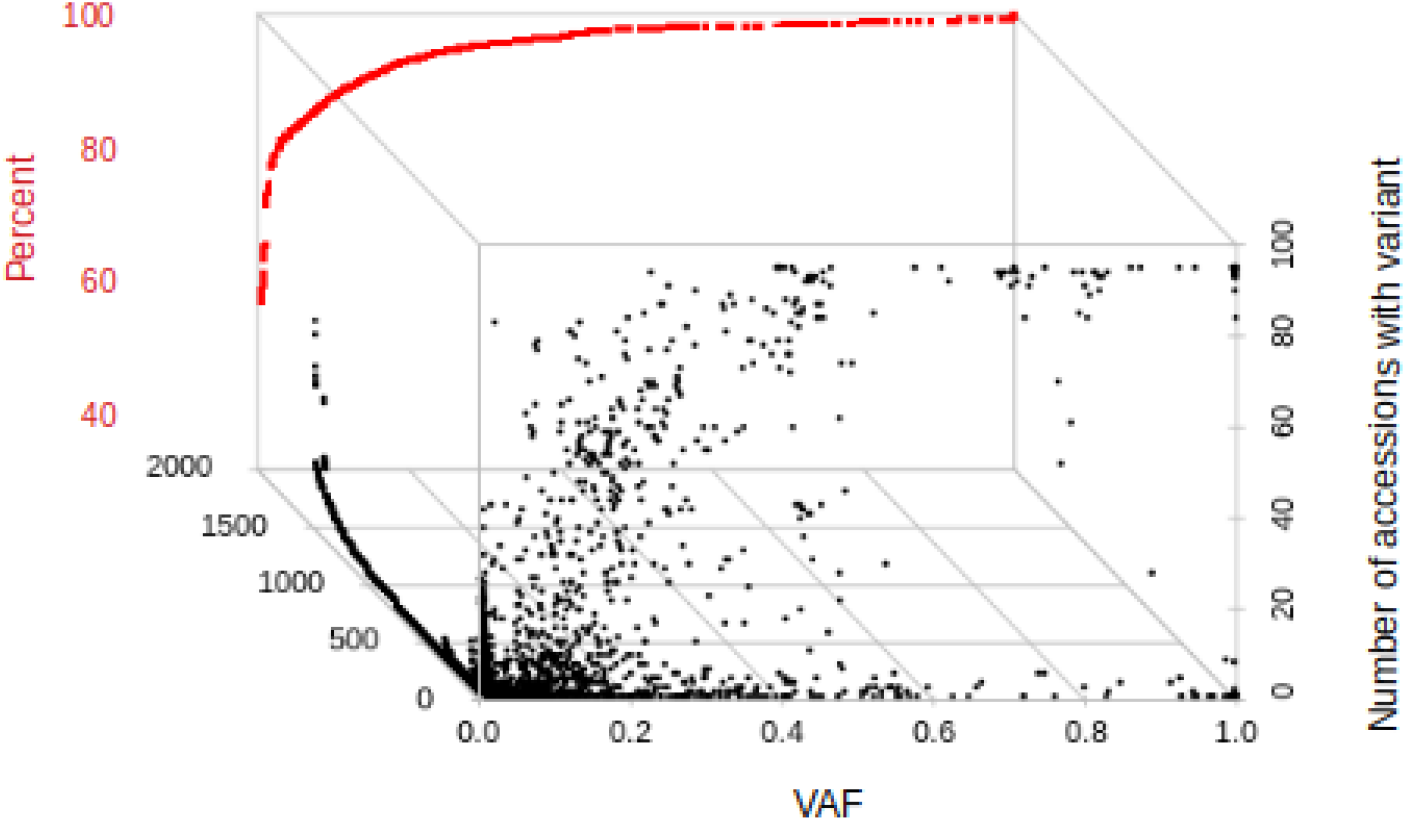
Scatter plot of Variant Allele Frequency (VAF) data from GATK HaplotypeCaller (black dots). VAF is plotted on the x-axis. Dots represent every predicted variant. The number of accessions predicted to harbor the variant is plotted on the y-axis. Data is plotted on the z-axis to separate different variants that share the same VAF and number of accessions. The percentage of the total data from VAF 0 to a specific frequency is overlaid in red. For example, 75% of all predicted nucleotide variants have a VAF of 0.05 or lower.

Variant calling was next carried out using SNVer and CRISP producing 1,570 and 3,261 variant calls, respectively. Similar to data produced with GATK, the highest percentage of variants are represented in the lowest VAFs (75% at 0.034 or lower for CRISP and 0.088 or lower for SNVer, Supplemental Figure 2). Private variants were also enriched at lower VAFs (Supplemental Figure 1). In total, 895 variants were common between the three methods (Figure 2). Within these common variants, the mean VAF and the number of accessions carrying the variant differed between the three algorithms used.

**Figure 2.**
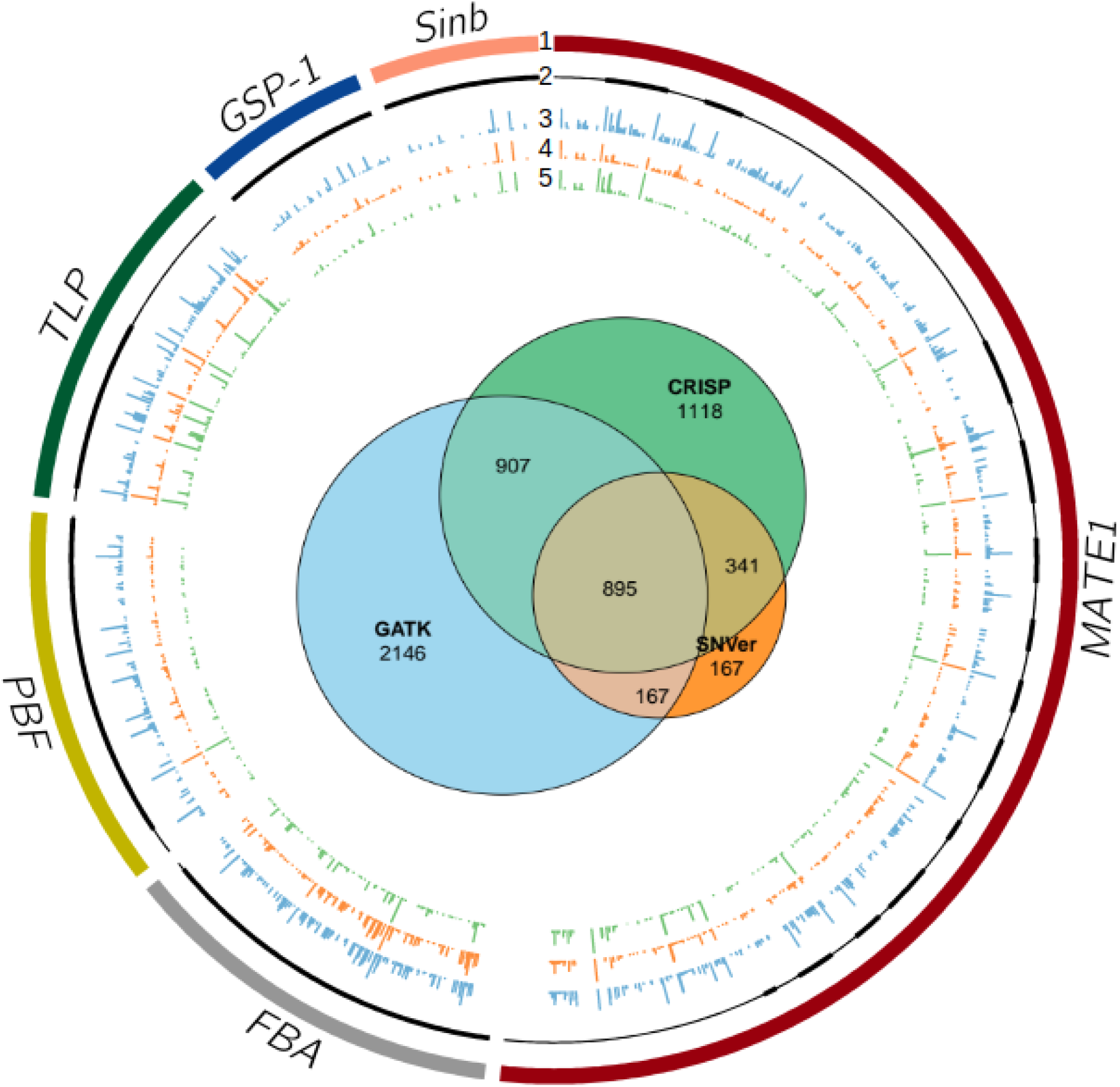
Common and unique variants called by GATK, SNVer and CRISP. The Venn diagram shows the overlap of variant calls for the three algorithms (interior image). Eight hundred and ninety-five variants were commonly identified. The outer image is a Circos plot of the common variants. Only the PCR amplified regions of gene targets are displayed (track 1). Gene models are shown on track 2 with exons and introns represented by thick and thin black lines, respectively. Tracks 3, 4, and 5 show the position and frequency (indicated by bar height) of variants predicted by GATK, SNVer and CRISP, respectively.

The effect on gene function of putative variants was evaluated with SNPeff and SIFT. This resulted in 695 putative deleterious variants from GATK, 171 from SNVer and 578 from CRISP, with 74 putative deleterious variants common to all three algorithms (Table 1, Supplemental Table 2, Supplemental Figure 3). Deleterious alleles with a high maximum VAF (the highest VAF reported in an accession) and present in only one accession were recovered along with alleles with a high maximum VAF that were present in 90 or more accessions (Supplemental Table 2). Alleles with a maximum VAF less than 0.4 were also identified, suggesting the presence of rare alleles segregating within an accession. In the GATK data set, for example, 29 of the 74 predicted deleterious common variants have a maximum VAF between 0.047 and 0.391 and are found in 1 to 21 accessions (Supplemental Table 2, Supplemental Figure 4).

**Table 1.** Missense, nonsense and silent changes with different variant calling methods.

|  | GATK | SNVer | CRISP | Common variants |
| --- | --- | --- | --- | --- |
| Missense | 1183 | 336 | 868 | 164 |
| Nonsense | 14 | 7 | 9 | 2 |
| Total | 1770 | 602 | 1322 | 348 |

Within target genes, 18 to 443 polymorphic positions were detected consistently by the three algorithms, corresponding to one SNP or InDel every eight to ten bp of sequence for five of the analyzed genes. For the sixth gene, *Sinb*, this frequency was markedly lower, with one SNP per 25 bp. The number of putatively deleterious variants per gene ranged from 11 (*GSP-1*) to 21 (*MATE1*), corresponding to one deleterious variant every 40 to 80 bp, with exception of *Sinb*, where only three deleterious variants were identified in 447 bp of coding sequence. Previous data on genic variation was available solely for *MATE1* (a total of 112 unique variants from 26 sequences deposited in GenBank as of 7^th^ January 2020) and Sinb with seven unique variants reported (Liu et al., 2017). The present study identified 62 new variants in *MATE1* coding sequence, including seven putatively deleterious, and 15 new variants in *Sinb*, including all three putatively deleterious variants. Most new variants identified in *MATE1* and *Sinb* were private or rare (median of the number of accessions with a given variant equaled two in *MATE1* and one in *Sinb)*.

The presence of predicted variants was assayed using Sanger sequencing of *Sinb* amplicons in a single individual plant from each of eight accessions. Twelve variants were predicted in this set. Only variants reported by all three algorithms for the tested accession, and where the lowest VAF was greater than 0.295 were validated (Supplemental Table 3). Because allele frequencies were calculated from a pooled DNA sample, it was concluded that lower frequency alleles likely represent alleles that are not present in every seed of an accession. Subsequent validation assays were carried out whereby multiple plants from each accession were assayed independently. In CAPS and Sanger sequencing assays on *MATE1, PBF*, and *Sinb* amplicons, eleven out of 16 tested variants were recovered when sampling between six and 16 plants (Table 2). Observed allele frequencies calculated from the number of plants harboring the tested sequence difference varied from the frequencies predicted from the amplicon sequencing data. Five variants had observed VAF closest to GATK predictions, three variants were closest with SNVer and three with CRISP. The four variants not recovered by CAPS or Sanger assays had frequencies reported by GATK below 0.15. Failure to recover low frequency alleles may have resulted from testing an insufficient number of individuals.

**Table 2.**
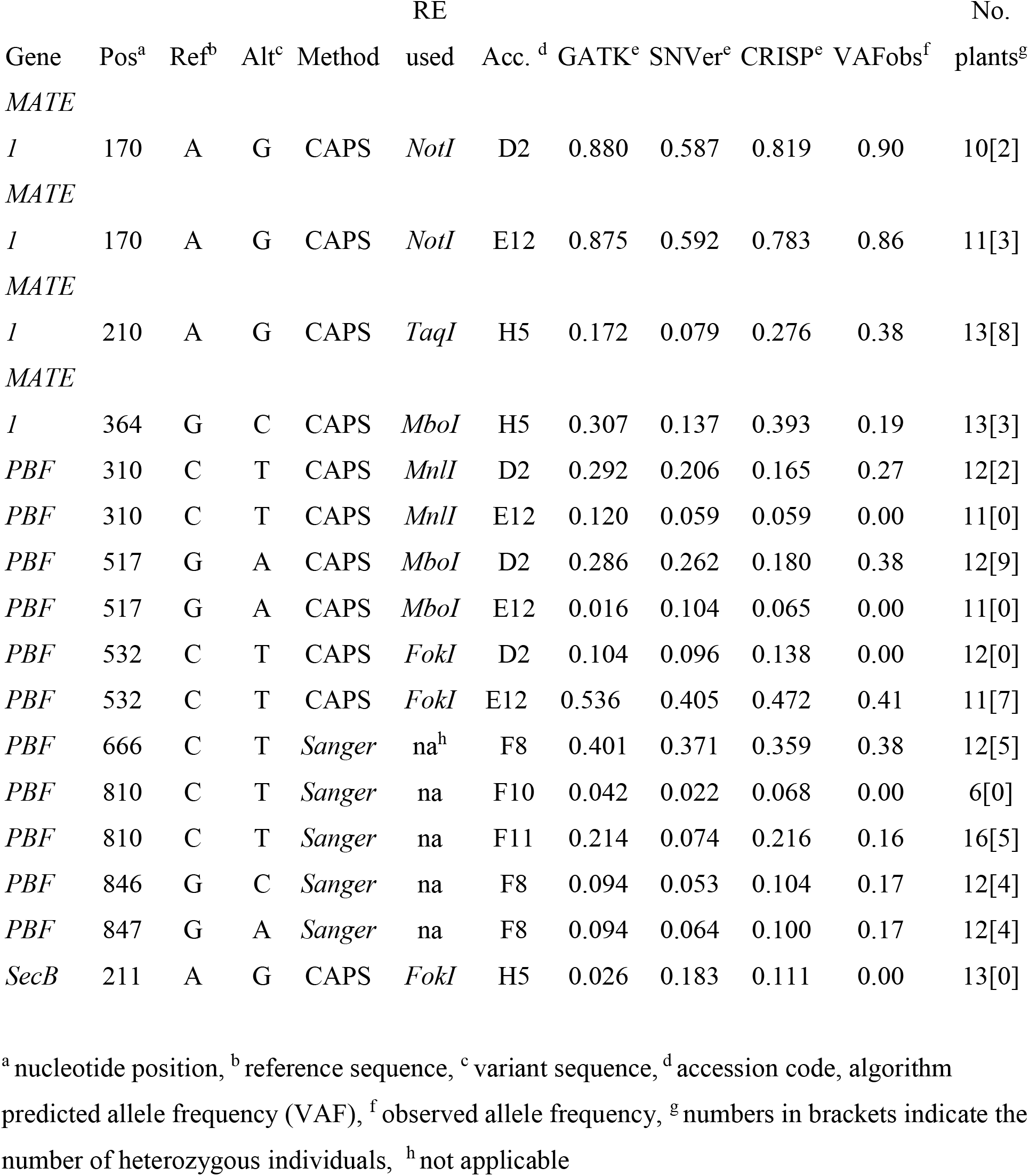
CAPS and Sanger validation of variants in multiple single plants of an accession.

### Phylogenetic relationships between populations and comparison of VAF distributions

The relationship between accessions was evaluated by creating a Neighbor Joining (NJ) tree based on Nei’s genetic distance (Figure 3). This resulted in accessions divided into six clusters (I-VI), with cluster I containing mostly cultivars, including the majority of modern cultivars analyzed. Nevertheless, a coincidence of the clustering with improvement status could not be observed. The accessions *S. sylvestre* (abbreviation B6 on Figure 3), *S. strictum subsp. kuprijanovii* (F8) and *S. strictum subsp africanum* (B7), were indicated as the most divergent of the analyzed set, which is in agreement with results of previous genome-wide analyzes of rye germplasm (Bolibok-Bragoszewska *et al*., 2014; Al-Beyroutiova *et al*., 2016; Rabanus-Wallace *et al*., 2019). Conversion of VAF values to genotyping scores was used to evaluate clustering. Different ranges of VAF were used to define heterozygous variants. This resulted in a changed clustering of the populations at each range tested (Supplemental Figure 5).

**Figure 3.**
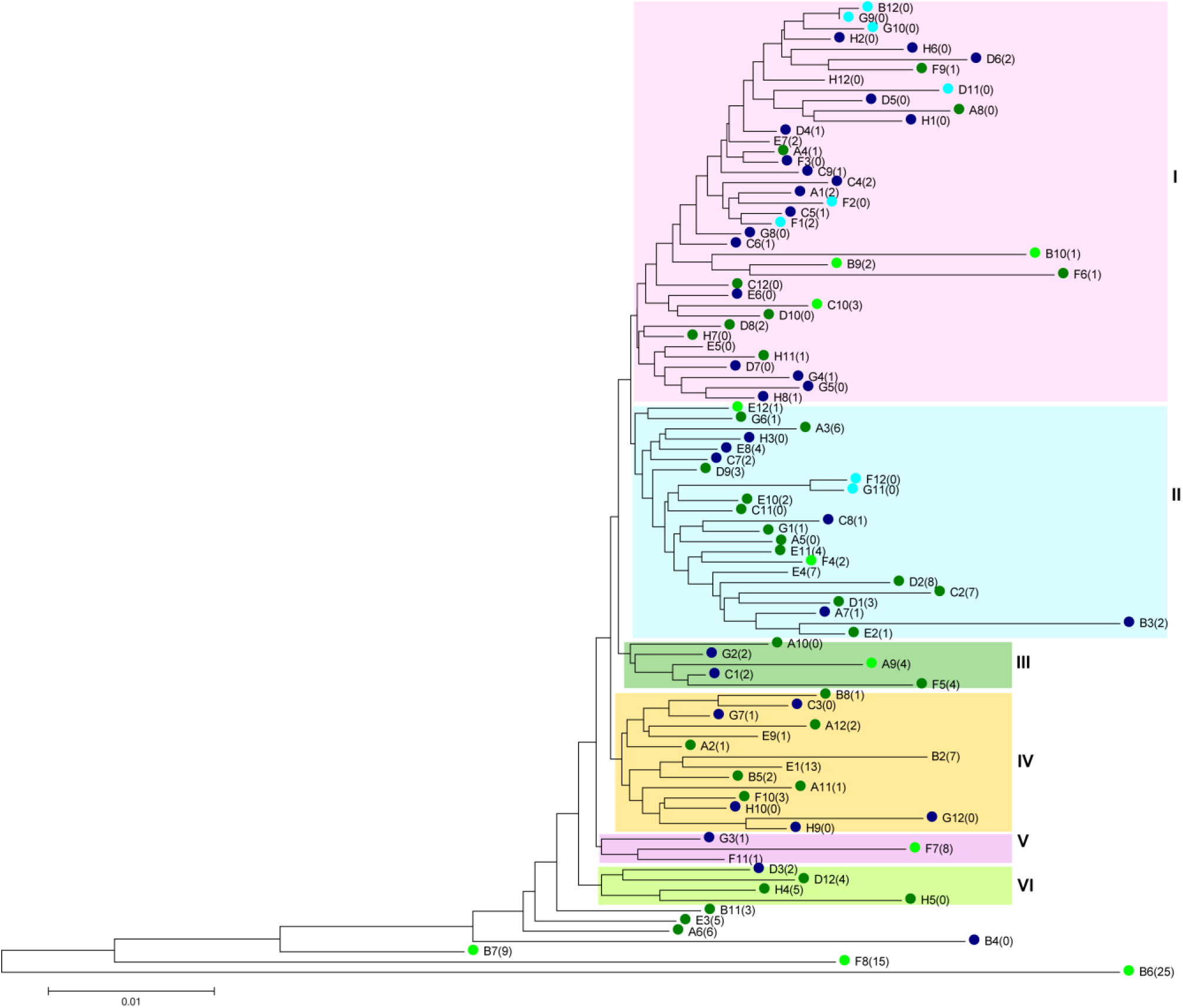
Neighbor Joining tree based on from Nei’s genetic distance calculated from VAF values reported by GATK for 895 variants detected in common by tree algorithms, showing genetic relationships between 95 rye accessions. To simplify the output, accessions are referred to by the 96 well plate coordinates, which are also included in the accession list (Supplementary Table 4). Numbers in brackets indicate private alleles identified in the respective accession. Colors indicate improvement status: light blue – modern cultivar, dark blue – historic cultivar, dark green – landrace, light green – wild accession.

Private variants occurred in all germplasm groups included in the study: modern cultivars, historic cultivars, landraces and wild accessions. In the group of commonly identified variants, the number of private variants per accession coincided with the domestication status: private variants were most frequent in wild accessions (five to seven per accession), followed by landraces with approximately two variants per accession, and historic and modern cultivars with less than one private variant per accession. Private variants in wild accessions also had the highest VAF values (mean 0.4-0,45, median 0.22-0.28). In the remaining germplasm groups mean VAF and median VAF did not exceed 0.12 and 0.007, respectively. The number of private variants varied from 0 (32 accessions) to 25 in *S. sylvestre* (B6) (Figure 3). Among the cultivated rye (*S. cereale* ssp. *cereale*) accessions, the highest number of private variants (7) was observed in a landrace from Bosnia and Herzegovina (C2) and also in a *S. cereale* ssp. *cereale* accession of unknown improvement status from Israel (E4).

VAF values were used to prepare violin plots in order to qualitatively compare accessions. Several distinct patterns of allele frequency distribution were observed (representative examples shown in Figure 4). Based on the results of two-part Wilcoxon test of pairwise comparisons of VAF distributions, rye accessions were grouped into 20 clusters ranging in size from 1 to 16 (Supplemental Figure 6). Five accessions, characterized by a high proportion of variants with high VAF values were consistently recognized as markedly different from the rest: *S. sylvestre* (B6), *S. strictum subsp. kuprijanovii* (F8), historic cultivars Imperial (B4), and Otello (G12) and landrace R1040 (F6) (Supplemental Table 4).

**Figure 4.**
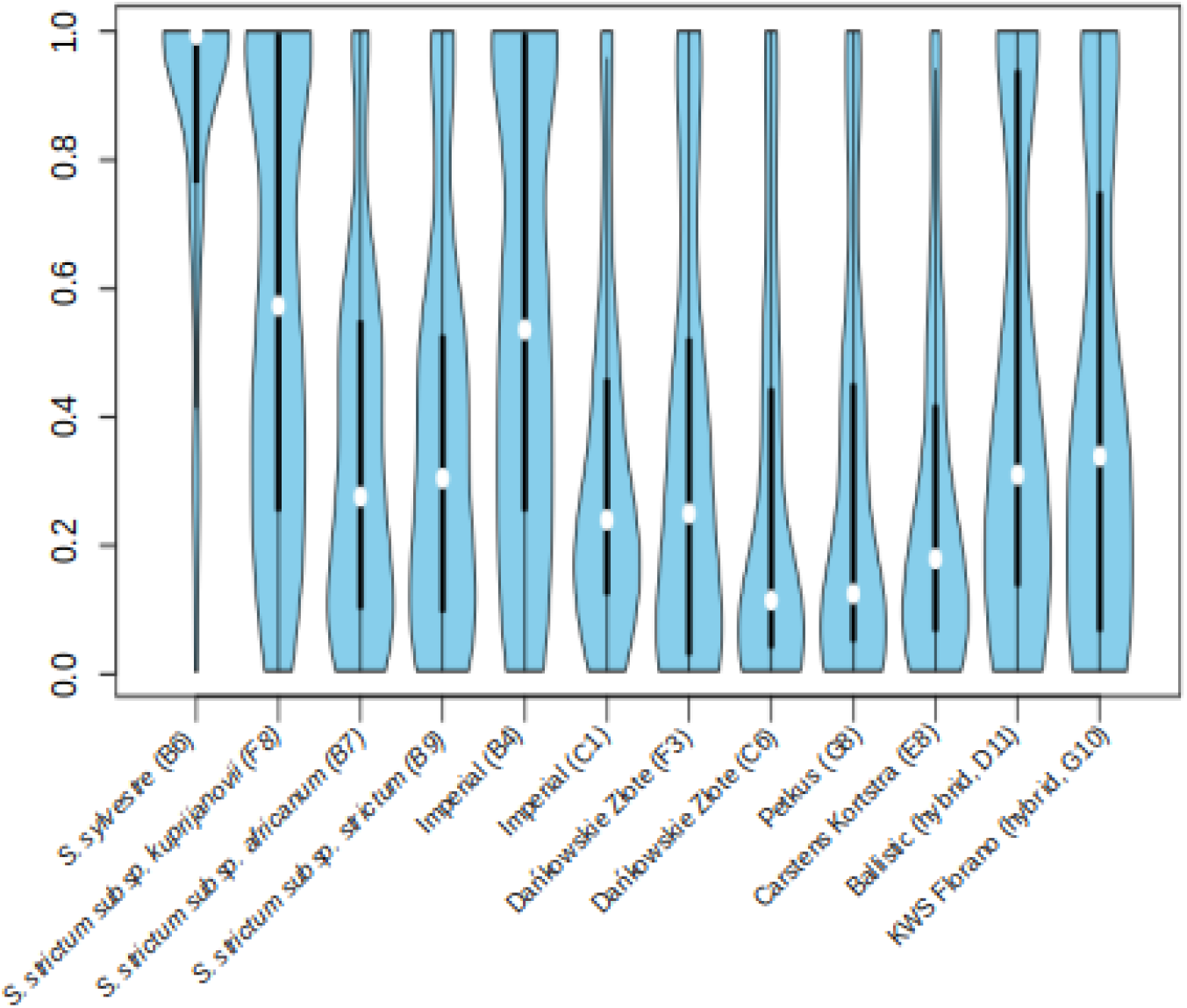
Violin plots of GATK VAF values from selected accessions for variants predicted in common by GATK HaplotypeCaller, SNVer and CRISP.

## DISCUSSION

To evaluate the distribution of frequencies of alleles within a landrace or cultivar, we chose to sample 96 plants from each accession of rye selected for our study. This allows the recovery of i) sequence differences compared to the reference sequence used that are fully homozygous (those with an allele frequency of 1), ii) heterozygous variants present in all pooled plants and iii) variants of lower frequencies that are not present in every seed in the seed stock used to represent an accession. To streamline the approach, tissue from each plant was collected and pooled prior to DNA extraction. The experiment was designed such that an allele found in a single plant could be identified. High coverage values were found in all DNA pools suggesting that each pool was suitable for PCR amplification and sequencing. Deviations in coverage values, therefore, likely resulted from differences associated with the quantification, normalization and pooling of PCR products. Such variations were recently reported in a study comparing tomato, cassava and barley amplicon sequencing data sets (Tramontano *et al*., 2019). The study revealed that minor coverage improvements could be achieved through the addition of extra quantification methods. Alternative approaches to increase read coverage at all nucleotide positions include increasing sequencing yields by adjusting the number of samples in an experiment and/or the number of target genes (amplicons) used in a single sequencing run. As sequencing costs drop, it may prove more cost effective and faster to simply produce more base pairs of data per experiment than to fine-tune the other experimental parameters.

Many applications employing next generation sequencing of genomic DNA involve the evaluation of sequence variations in diploid samples. Even in optimized diploid conditions, a balance is struck between maximizing allele calling sensitivity to reduce false negative errors and reducing the sensitivity in order to lower false positive errors. For example, when using GATK HaplotypeCaller with settings for diploid samples, Li et al reported that more than 80% of false positive errors in diploid rice were at an allele frequency below 40% (Li *et al*., 2019). When sequencing non-pooled samples, setting an allele frequency threshold of >40% for heterozygous variants therefore reduced false positive errors. Such optimizations are more challenging in pooled samples. Algorithms such as GATK HaplotypeCaller, SNVer and CRISP provide parameter settings to call low frequency variants. Yet, optimal parameters still need to be determined. For example, evaluation of six SNP calling algorithms in tomato TILLING samples pooled either 64 or 96 fold revealed that accuracy ranged between 89.33 and 25.33 % when comparing to Sanger validated SNP mutations (Gupta *et al*., 2017). That work described technical differences between different algorithms and concluded that accuracy is improved when a variant call is predicted by at least two algorithms. In cassava, up to 281 different accessions were pooled together prior to sequencing in an approach designed to quickly identify putative deleterious alleles (Duitama *et al*., 2017). In that study 24% (79/325) of called variants were predicted by four algorithms tested.

The experimental design for rye differed from previous studies in order to allow the discovery and analysis of intra-accession allele variation. The assay was designed to recover two types of what can be considered “rare” variation. The first type of rare variants are alleles that are found in only one accession (known as private alleles) or very few accessions in the tested set, and occur with a high frequency within the respective accessions. This type of rare variation is easily recovered using conventional genotyping and resequencing as alleles can be recovered through assay of a single seed (Wang *et al*., 2018; Balfourier *et al*., 2019) The second type of rare variants are more difficult to discover. These variants segregate at a low frequency within an accession and are never found at high frequency in any tested accession. To recover this type of rare variant requires the sampling of multiple individual seed per accession. As such, these alleles are hidden from discovery when using traditional methods that sample one or few seed per accession. Using pooled amplicon sequencing we have recovered both types of rare alleles in the tested rye accessions. Importantly, the presence of variants that segregate at a low frequency within an accession, and are never found at high frequency in any tested accession, suggest that a broader genetic diversity can exist in germplasm collections than previously known. We expect this to be most common in outcrossing species like rye where admixtures of alleles are frequent.

Variants with mean VAF between 0.7 and 1 represented between 1.87 and 3.06% of all predicted alleles, depending on the algorithm used. In this set of variants, between 41 and 49% are private alleles found in only one accession (Figure 1, Supplemental Figures 1 &2). The highest number of variants were found in the lowest VAFs. It is expected that false positive errors will increase as the number and percentage of reads supporting the alternative allele decreases. Studies have been carried out on errors associated with MiSeq paired end sequencing, but a thorough investigation into errors in pooled samples has not been reported (Schirmer *et al.*, 2015). False positive errors are expected to be random and therefore infrequently independently predicted when applying multiple variant calling algorithms. Indeed, of the 895 variants common to GATK, SNVer and CRISP, only 20% had a predicted mean VAF of 0.038 or lower, a reduction of more than 50% from the data from any single algorithm. Further experiments are required to determine what, if any, percentage of the sub 0.038 VAF variants predicted by all three algorithms are false positive errors. This requires extensive genotyping, as many individual seed need to be tested to ensure true variants are recovered. In the present study, genotyping assays using approximately 10 seed per accession were sufficient to validate alleles with a VAF of 0.15 or higher that were predicted in the same accession by all three algorithms. We expect it is necessary to test more than 100 seeds per accession to validate the lowest frequency alleles in the data. Some very low frequency false positive errors are expected and may result from biological contamination, for example, from pollen contamination on the leaf tissue collected. This can be ruled out in the present study because seedlings were grown, and tissue collected in growth room conditions where there were no rye plants flowering. Sample to sample cross contamination of DNA or PCR product may also be a source of low VAF false positive errors. Sixty-five percent of sub 0.038 VAF variants commonly predicted by all algorithms were found in more than one accession. However, 96 % had a maximum predicted VAF of less than 10%, and the highest maximum VAF was 24.5%. This means that a large volume of accidental liquid transfer between samples would be needed to create a detectable false positive. With the caveat of possible very low frequency false positive errors, we conclude that selecting variants commonly called by multiple algorithms may reduce errors and serves as a useful method to prioritize alleles for further study.

We found qualitative evaluation of VAF values using violin plots to be useful to estimate the influence of a taxon’s reproductive biology, preservation history and breeding on the genetic composition of an accession. For example, one of the outlier accessions identified is this study is *S. sylvestre* (B6). Molecular marker-based analyses of genetic diversity indicated this self-pollinating taxon as the most divergent in genus *Secale* (Bolibok-Bragoszewska *et al.*, 2014; Al-Beyroutiova *et al.*, 2016; Schreiber *et al.*, 2018). Its large proportion of high VAF variants (Figure 4) likely corresponds to homozygosity for alternative alleles, since reference sequences used during variant calling originated from cultivated rye accessions. Another outlier, *S. strictum* subsp. *kuprijanovii* (F8), is a perennial outbreeder, also genetically divergent from *S. cereale*. However, its violin plot differs markedly from plots obtained for the other two *S. strictum* samples included in the study, *S. strictum* subsp. *africanum* (B7) and *S. strictum* subsp. *strictum* (B9), which might indicate a sample tracking mistake during genebank preservation or laboratory handling, or a bottleneck during preservation. A sample of Imperial cultivar (B4), widely used in cytological studies, originating from the collection of A. J. Lukaszewski (UCLA, Riverside) showed an approximately equal abundancy of variants with all possible VAF values and differed clearly from another sample of Imperial, (C1), obtained from IPK Gatersleben genebank. Less pronounced (although also statistically significant) differences were also observed between the two samples of cultivar Dankowskie Zlote (C6 and F3), obtained from different sources. Samples of hybrid cultivars from KWS, such as Ballistic (D11) and KWS Florano (G10), exhibited a higher percentage of VAF values in the range 0.3-0.5, with median ca. 0.3, and also higher percentage of AF values close to 1.0, in comparison to population cultivars included in the study, such as Dankowskie Zlote (F3 and C6), Petkus (G8), or Carstens Kortstra (E8), which is consistent with the use of the three line system in the development of hybrid rye cultivars.

In this study we analyzed six genes linked to biotic and abiotic stress resistance and seed quality. Using deep sampling and pooled amplicon sequencing numerous new variants were identified, (including putatively deleterious ones), in each of the analyzed genes, providing potential targets for future functional studies and, eventually, inclusion in breeding schemes in rye and related species (wheat, triticale). Consistent with a high diversity of the germplasm set used (with respect to domestication status and origin) we obtained a several fold higher estimate of SNP frequency in rye (on average one SNP or InDel every 12 bp), than those reported in the past: 1 SNP/52bp (Li *et al.*, 2011), 1 SNP/58bp (Varshney *et al.*, 2007) or 1 SNP or InDel/31 bp (Bauer *et al.*, 2017). In agreement with the results of previous genome-wide, DArT-marker based characterization of genetic diversity in rye (Bolibok-Bragoszewska *et al.*, 2014), data obtained in the present study on distribution of private alleles among germplasm groups indicates that the genetic diversity in modern rye cultivars is relatively narrow, with less than one private allele identified per modern cultivar tested, and provides further evidence for the value of rye PGR in genetic research and crop improvement, with more than five private alleles identified per accession, stressing the importance of conservation and characterization efforts. On the other hand, the clustering of the accessions in the NJ tree generated based on the VAF of 895 variants detected in common did not agree with the improvement status of the accessions, suggesting, that selective pressures other than breeding practices have influenced the diversity of the genes analyzed.

This study also points out that, in case of open pollinated populations (due to the high within-accession variability), the sampling of a single individual or a small number of individuals from an accession most likely results in an inaccurate and perhaps even misleading representation of genetic relationships between the accessions. This can be seen in NJ trees produced based on conversion of VAF values into genotyping-like scores, where a different clustering of accessions was observed at each range of VAF used to define heterozygous variants (Supplementary Figure 5).

The approach of deep sampling and pooled amplicon sequencing allows discovery of variants in candidate genes and also an evaluation of the effect of variants on gene function. This provides an additional filter to prioritize variants. The SIFT program was used to identify 73 putative deleterious alleles commonly identified by the three variant calling algorithms. This data set contained different classes of alleles for example, homozygous variants found in one or few individual accessions (private deleterious alleles, the first category of rare variants described above). Homozygous variants present in more than 90 accessions were also recovered. Interestingly, putative deleterious variants were also identified where the maximum VAF was between 0.15 and 0.3 and the variant was found in only one or two individual accessions (the second category of rare variant). This suggests that alleles are segregating within rye accessions at low fractions that may affect gene function and potentially plant phenotype. Such variants would go undiscovered in conventional GBS or WGS assays where only one or two seed per accession are sampled, and may be useful for functional genomic characterizations and breeding. Further studies are being designed to evaluate the different classes of putative deleterious alleles. For example, homozygous private alleles may represent alleles where a fitness penalty results in the allele having been expunged from most populations. Homozygous putative deleterious alleles present in most tested accessions may represent alleles with no fitness penalty, or may represent alleles that have no negative effect on fitness under their natural growing conditions (e.g. low aluminium in the soil). Possible mechanisms for the maintenance of rare low frequency alleles in populations, including meiotic effects, can also be investigated.

The rye amplicons used in this study were generated before the release of the rye genome (Rabanus-Wallace *et al.*, 2019). It is expected that the recent release of the rye reference genome will enable improvements in gene target selection and primer design. Reference genomes have been produced for few of the hundreds of thousands of plant species existing on the planet. Because pooled amplicon sequencing does not require complete genome sequence, we expect that the approach described for rye can be adapted for many plant species and can facilitate better characterization of existing rich germplasm collections. We predict that flexible and low-cost methods for recovery of rare genetic variation will support future efforts to promote sustainable food security.

## MATERIALS AND METHODS

### Plant material

Ninety-five accessions of rye, each represented by a pooled sample comprising DNA of 96 individual plants, were analyzed in the study. This set included 90 accessions of *S. cereale*, among them 8 modern cultivars, 34 historic cultivars, 35 landraces, and 5 accessions of other *Secale* taxa, representing various geographic regions. In total 10 accessions from this set were described as wild/weedy. Seeds were obtained from several sources including genebanks and breeding companies (Supplemental Table 4).

### Genomic DNA extraction, quantification and pooling

Seeds were placed in a growth room in containers lined with moist paper towels. Ten days after germination a 20 mm long leaf segment was harvested from each plant. For each accession 96 plants were sampled. Leaf segments from 16 plants of the same accession were collected into one 2 mL centrifuge tube, with six tubes from 16 individual plants obtained for each accession. Collected leaves were freeze-dried in an Alpha 2-4 LDplus lyophilizator (Christ), for 18h at −60 °C, 0.011 mbar, followed by 1h at −64°C, 0,006 mbar and ground to fine powder using a laboratory mill MM 301 (Retsch) for 2.5-5 min at frequency 30.0 1/s. Genomic DNA was extracted using Mag-Bind Plant DNA DS Kit (OMEGA Bio-Tek) following manufacturer’s protocol. Quality and quantity of DNA was assessed using spectrophotometry (NanoDrop2000, Thermo) and electrophoresis in 1% agarose gels stained with ethidium bromide. DNA concentration of each sample tube was adjusted to 100 ng and an equal volume of all samples from an accession were pooled together.

### Primer design and PCR amplification of target genes

Sequences of six target genes: aluminium activated citrate transporter (*MATE1*, also known as *AACT1*), taumatin-like protein (*TLP*), fructose-biphosphate aldolase (*FBA*), prolamine-box binding factor (*PBF*), secaloindoline-b (*Sinb*) and grain softness protein *(GSP-1*) were retrieved from GenBank (Supplemental Figure 7, Supplemental Table 5). The entire sequences of *Secb* and *GSP-1* genes (456 and 506 bp, respectively) were amplified using primers described, respectively, by Simeone and Lafiandra (Simeone and Lafiandra, 2005) and Massa et al. (Massa *et al.*, 2004). For genes *FBA*, *MATE1*, *PBF* and *TLP* primer pairs for generation of overlapping, ca. 600 bp long amplicons, covering the entire gene sequence were designed using Primer-BLAST (Ye *et al.*, 2012). Primer pairs were tested using the DNA of rye inbred line L318 and those producing single product of expected length were used for amplification of gene fragments from pooled DNAs. Primer design and all other assays described in this work were carried out before the public release of the rye genome. PCR set up was as follows: 200 ng of template DNA, 2.5 mM MgCl2, 0.2μM of each primer, 0.2 mM of each dNTP, 1x Dream *Taq* Green buffer, 0.5 U Dream *Taq* DNA polymerase (Thermo Scientific). The reactions were carried out in 25 μL in Mastercycler epgradient S (Eppendorf) thermal cyclers. For all primer pairs the thermal profile of initial denaturation for 60s at 95°C, 30 cycles of 30s at 95°C, 30s at 56°C and 60s at 72°C, followed by final extension for 5 min at 72°C was used. A volume of 5 μL from each reaction was used to check the amplification success using electrophoretic separation in 1.5% agarose gels stained with ethidium bromide. PCR products were shipped to Plant Breeding and Genetics Laboratory, Joint FAO/IAEA Division, International Atomic Energy Agency (Seibersdorf, Austria) for further processing.

### PCR product quantification and pooling

PCR products were quantified using egel 96well gels (Thermo Fisher Scientific) and quantitative lambda DNA standards as previously described (Huynh et al., 2016). PCR product concentration was adjusted to 10 ng/ul in TE. All PCR products from a single gDNA pool were then pooled together. Pooled PCR products from each of the 95 accessions were then quantified using the Advanced Analytical^®^ Fragment Analyzer™ with the low sensitivity 1kb separation matrix with 30 cm capillaries (Advanced Analytical®#DNF935). All sample pools were normalized to 30 nM concentration in TE prior to library preparation.

### Library preparation and sequencing

Indexed DNA library for NGS was prepared using the TruSeq® Nano DNA HT Library Preparation Kit (Illumina, cat. 20015965) according to manufacturer’s recommendation. Indexed libraries were then quantified using a Q-bit fluorometer (Thermo Fisher Scientific) and pooled together at an equal concentration. The pooled library was diluted to 18 pM concentration. Sequencing was performed on an Illumina MiSeq® using 2×300 PE chemistry according to manufacturer’s protocol. The reads were de-multiplexed with the MiSeq Reporter software and were stored as FASTQ files for downstream analysis.

### Sequence evaluation

FASTQ files were aligned to target amplicons using BWA mem with commands -M -t 16 (Li and Durbin, 2009). Amplicon fragments were given target names that were used throughout the NGS analysis (Supplemental Table 5). Samtools view was used to convert from SAM to BAM format (Li *et al.*, 2009). Bam files are available in NCBI BioProject PRJNA593253 Coverage statistics were prepared using qualimap v.2.2.1-dev (García-Alcalde *et al.*, 2012). Variant calling was performed using three algorithms CRISP, GATK and SNVer. Parameters used for CRISP were –OPE 0, --poolsize 192 and –qvoffset 33 (Bansal, 2010). The GUI of SNVer was used with the following parameters: -bq20,-mq17,-s0,-f0,-pbonferroni=0.1,-a0,-u30, -n192,-t0 (Wei *et al.*, 2011). HaplotypeCaller (GATK 4.0) was used following best practices with default settings with the exception that ploidy was set to 192 (Poplin *et al.*, 2017). For each method, VCF files from individual pools were merged using bcftools. Following this, read group information was unified between the three files using picard tools AddOrReplaceReadGroups function (http://broadinstitute.github.io/picard/index.html). Data for calculation of allele frequency from the VCF files (called VAF in this manuscript) was extracted for each variant and each accession using R libraries vcfR (Knaus and Grünwald, 2017) and VariantAnnotation (Obenchain *et al.*, 2014) and used to produce AF tables. The potential effect of nucleotide variation on gene function was evaluated with SNPeff (Cingolani *et al.*, 2012). For this, a genome database was prepared using the build - genbank function. The effect of reported nucleotide variation was also evaluated with SIFT4G using a self-prepared genomic database with the fasta file of amplicon sequences used for mapping with BWA mem, a self-prepared gtf file and the uniref90 protein database (Vaser *et al.*, 2016). Venn diagrams were produced using the R package eulerr (https://github.com/jolars/eulerr).

### Evaluation of VAF distributions

Violin plots were drawn using the geom._violin function of ggplot2 in R (Wickham, 2016) on VAF values reported by GATK for the variants detected in common by three algorithms. VAF value distributions were compared pairwise using the two-part Wilcoxon test (Gleiss *et al.*, 2015) resulting in a pairwise matrix of 0s and 1s, with 1 indicating that for the given pair of populations the distributions of VAF values are different at a = 0.05. This matrix was then used for hierarchical clustering analysis with the haclust function of the R package stats.

### Evaluation of phylogenetic relationships between accessions

For the purpose of illustrating the relationships between rye populations analyzed, a Nei’s genetic distance (Tateno *et al.*, 1982) matrix was calculated using POPTREEW (Takezaki *et al.*, 2014) using VAF values reported by GATK for the variants detected in common by three algorithms and imported into MEGA 5. 2 (Tamura *et al.*, 2011) to produce a Neighbor Joining dendrogram. To simulate the effect of treating the accessions as individuals on the clustering, VAF value tables were converted to genotyping scores (with “0” meaning a reference allele homozygote”, “1” meaning a variant allele homozygote, and 2 meaning a heterozygote). Three settings were applied that use different VAFs to define heterozygous variants: i)VAF < 0.3 = 0; VAF ≥ 0.7 = 1; and values in between (greater than 0.3 and less than 0.7) = 2, ii) VAF < 0.4 = 0; VAF ≥ 0.6 = 1; and values in between = 2, and iii) VAF < 0.2 = 0; VAF ≥ 0.8 = 1; in between = The obtained genotype scores were used as input to GenAlEx 6.5 for calculation of Euclidean distances. Neighbor Joining trees were produced from the resulting distance matrices using MEGA 5.2.

### Validation of nucleotide variants

For validation of nucleotide variants CAPS assays were developed based on output of PARSESNP (Taylor and Greene, 2003), which provides a list of restriction endonuclease sites that are gained or lost due to the predicted SNV or indel. Serial Cloner 2.6.1. (http://serialbasics.free.fr/Serial_Cloner.html) software was used to digest *in silico* the gene fragment of interest and predict restriction patterns for reference and mutant alleles. New batches of seeds were sown for selected accessions. Tissue harvest, DNA isolation and PCR reaction were done separately for each plant, using the procedures described above. Restriction digestion was done for 20 minutes using 10 μL of PCR reaction as template and 1 μL of the restriction enzyme in the total volume of 20 μL. FastDigest restriction enzymes (ThermoFisher) with dedicated buffers were used. The digestion products were separated in 6% denaturing polyacrylamide gels (if the predicted products were shorter than 200 bp or differed in length by less than 50 bp) and visualized by silver staining as described by Targońska et al. (Targońska *et al.*, 2016), or in 1.5% agarose gels containing ethidium bromide. For Sanger sequencing-based validation of variants PCR reactions were sent to an external service provider. The analyzed plants were classified based on electrophoretic separation patterns/chromatograms as homozygous reference (RefRef), heterozygous (RefAlt), or homozygous variant (AltAlt). The variant frequency was calculated using the formula (RefAlt × 1 + Alt/Alt × 2)/ n × 2, where n is the total number of individuals analyzed.

## Author Contributions

Conceptualization, B.J.T. and H.B-B.; Methodology, A.H., J.J-C., B.J.T. and H.B-B.; Investigation, A.H., L.B., K.T., E.B., J.J-C., P.G., A.K., B.J.T., and H.B-B.; Writing – Original Draft, A.H., B.J.T. and H.B-B.; Writing – Review & Editing, A.H., J.J-C., B.J.T., and H.B-B.; Funding Acquisition, B.J.T. and H.B-B.; Resources, B.J.T. and H.B-B.; Supervision, B.J.T. and H.B-B.

## Acknowledgments

This research was funded by the Polish National Science Center grant No. DEC-2014/14/E/ NZ9/00285. Funding for DNA sequencing was provided by the Food and Agriculture Organization of the United Nations and the International Atomic Energy Agency through their Joint FAO/IAEA Program of Nuclear Techniques in Food and Agriculture. The authors declare that there is no conflict of interest.

## Supporting Information

**Supplemental Figure 1.** Percentage of private alleles (found in only one of the tested accessions) plotted by variant allele frequency (VAF). Data from GATK is plotted in light blue, CRISP in green and SNVer in orange.

**Supplemental Figure 2.** Scatter plots of variant allele frequency (VAF) data (black dots). VAF is plotted on the x-axis. Dots represent every predicted variant. The number of accessions predicted to harbor the variant is plotted on the y-axis. Data is plotted on the z-axis to separate different variants that share the same VAF and number of accessions. The percentage of the total data from VAF 0 to a specific frequency is overlaid in red. Variants predicted by CRISP are plotted in panel A, and by SNVer in panel B.

**Supplemental Figure 3.** Venn diagram of variants called by GATK, SNVer and CRISP predicted to be deleterious using SIFT.

**Supplemental Figure 4.** Lollipop chart of allele frequencies of GATK variants predicted deleterious by SIFT and also called by SNVer and CRISP. Each variant is assigned an arbitrary number (x axis) with maximum allele frequency values calculated from GATK VCF data is plotted on the y axis. Data is sorted into 5 distinct groups based on the number of accessions harboring the variant. This sorting is indicated by the colored ball at the end of the bar. Allele frequencies below 0.039 are not plotted.

**Supplemental Figure 5.** NJ dendrograms based on conversion of VAF values reported by GATK for variants identified in common into genotype scores (“0” = reference allele homozygote”, “1” = variant allele homozygote, “2” = heterozygote) using the following settings: A) i)VAF < 0.3 = 0; VAF ≥ 0.7 = 1; and values in between (greater than 0.3 and less than 0.7) = 2, ii) VAF < 0.4 = 0; VAF ≥ 0.6 = 1; and values in between = 2, and iii) VAF < 0.2 = 0; VAF ≥ 0.8 = 1; in between = 2. Colors of the nodes correspond to colors of the clusters in the NJ dendrogram derived from VAF data (Figure 3, main manuscript) and indicate membership of the respective accessions in the clusters of the NJ dendrogram derived from VAF data.

**Supplemental Figure 6.** Dendrogram showing relationships between distributions of VAF values (shown as violin plots) for 95 accessions (based on results of the two-part Wilcoxon test).

**Supplemental Figure 7.** Target regions used in this study. Introns are colored blue, non-coding sequence grey and exons green. Relative nucleotide positions in base pairs are listed.

**Supplemental Table 1.** Sequencing coverage for each nucleotide position in the experiment.

**Supplemental Table 2.** Allele frequencies and number of accessions harboring alleles of predicted deleterious variants common to GATK, SNVer and CRISP.

**Supplemental Table 3.** Sanger sequencing validation of variants in single plants from an accession.

**Supplemental Table 4.** Accessions used in this study.

**Supplemental Table 5.** Primer sequences used in this study.

## Notes

### Competing Interest Statement

The authors have declared no competing interest.

### Summary of Updates

Abstract and manuscript text revised Figures updated Author emails updated Supplemental files updated

