## Supplementary figures and images for "Deep sampling and pooled amplicon sequencing reveals hidden genic variation in heterogeneous rye accessions"

### Supplemental Figure 1

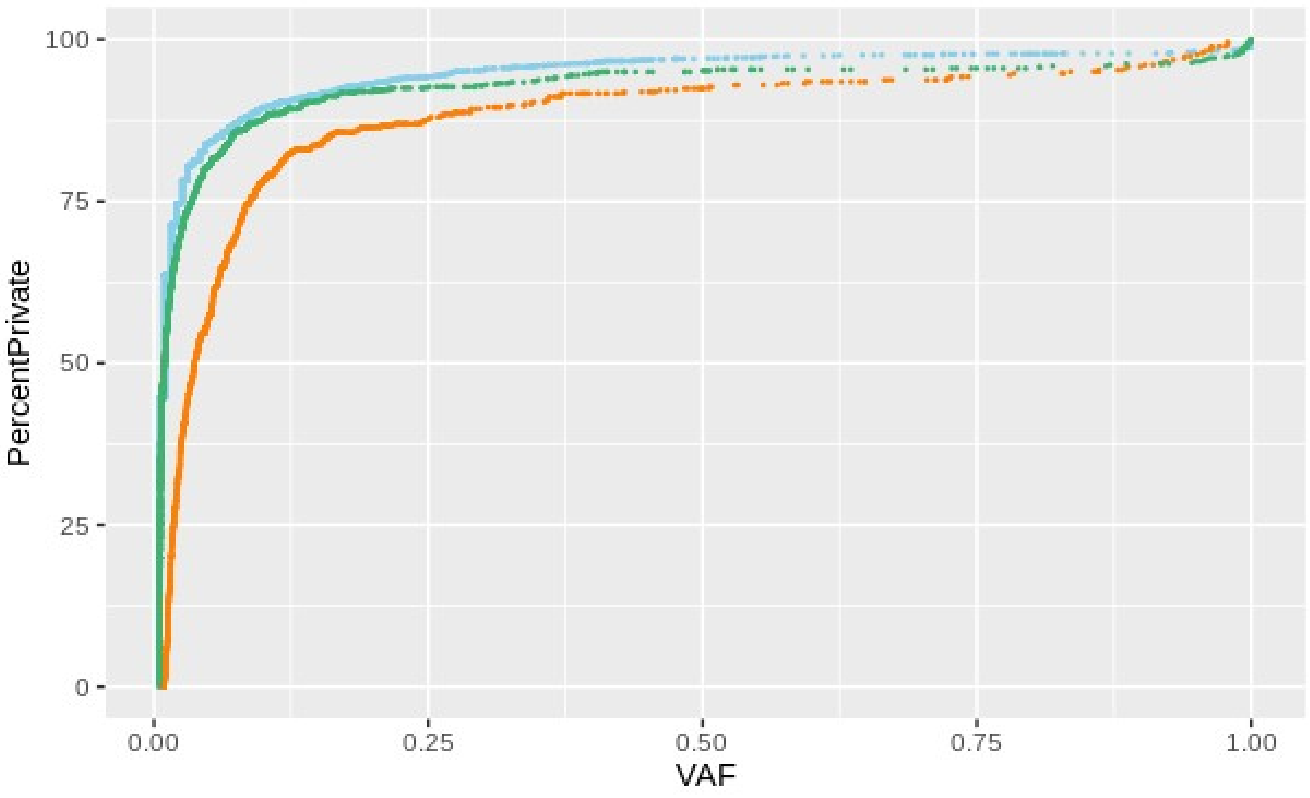

### Supplemental Figure 2

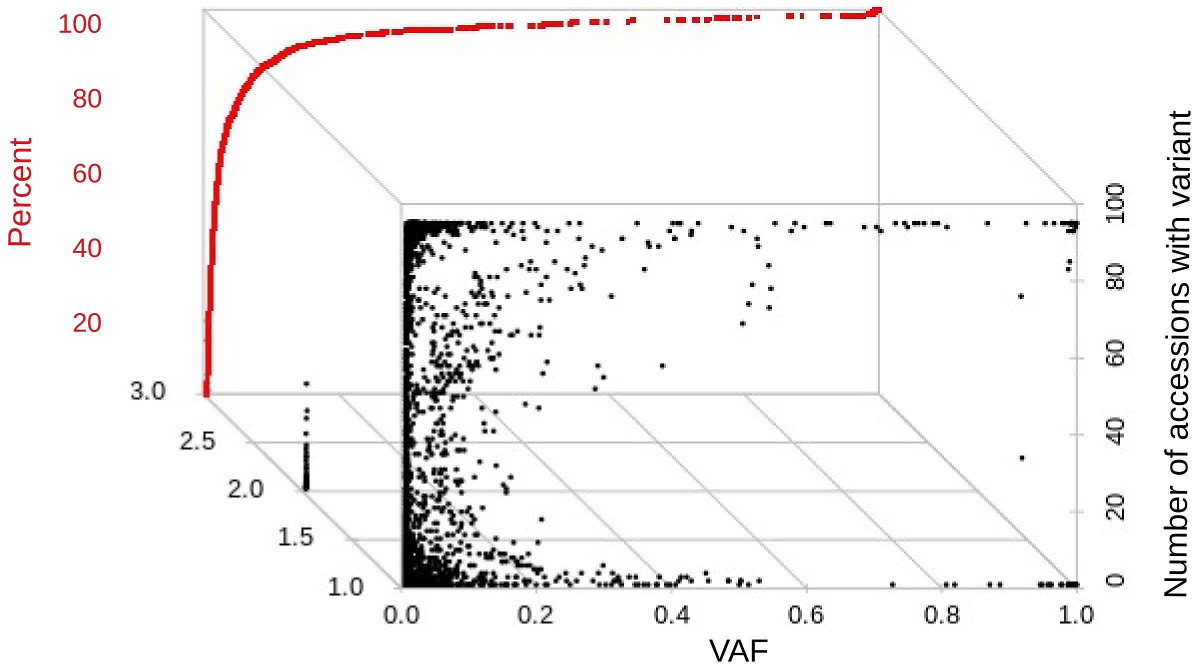

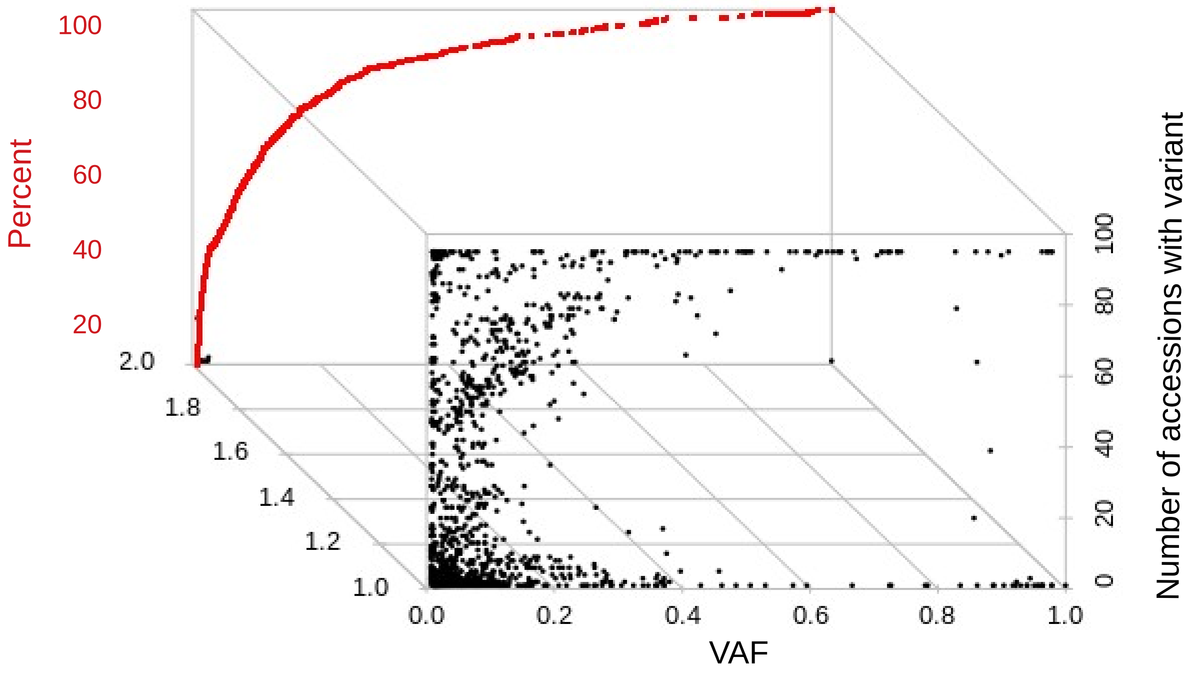

### Supplemental Figure 3

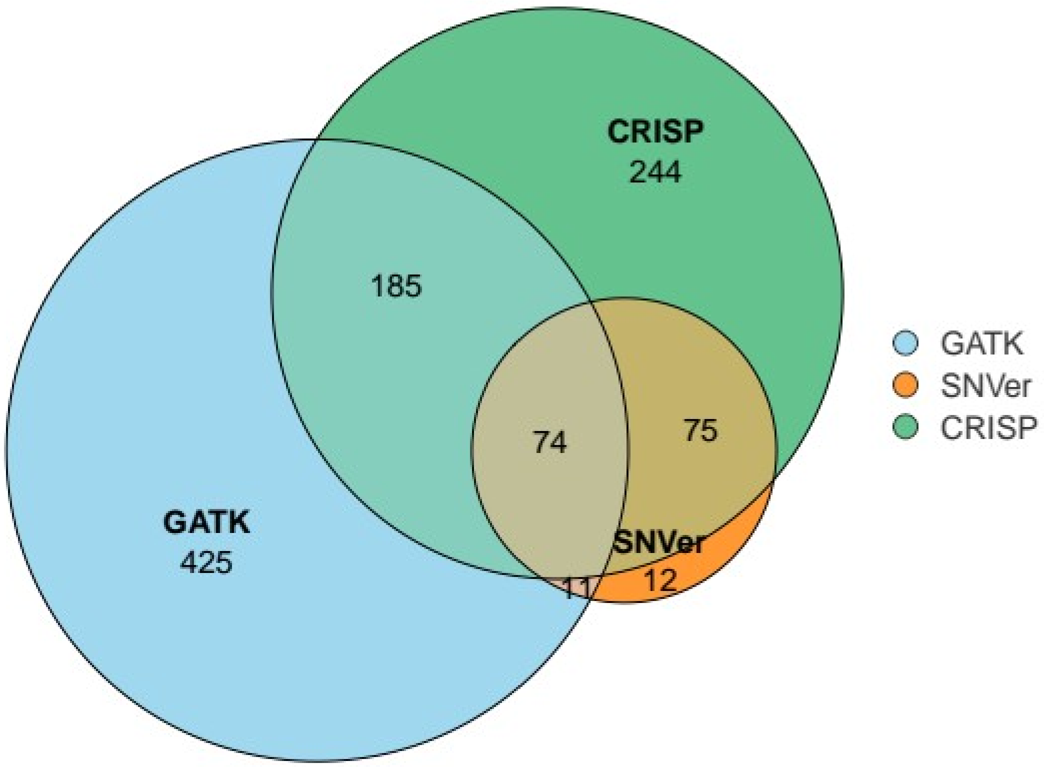

### Supplemental Figure 4

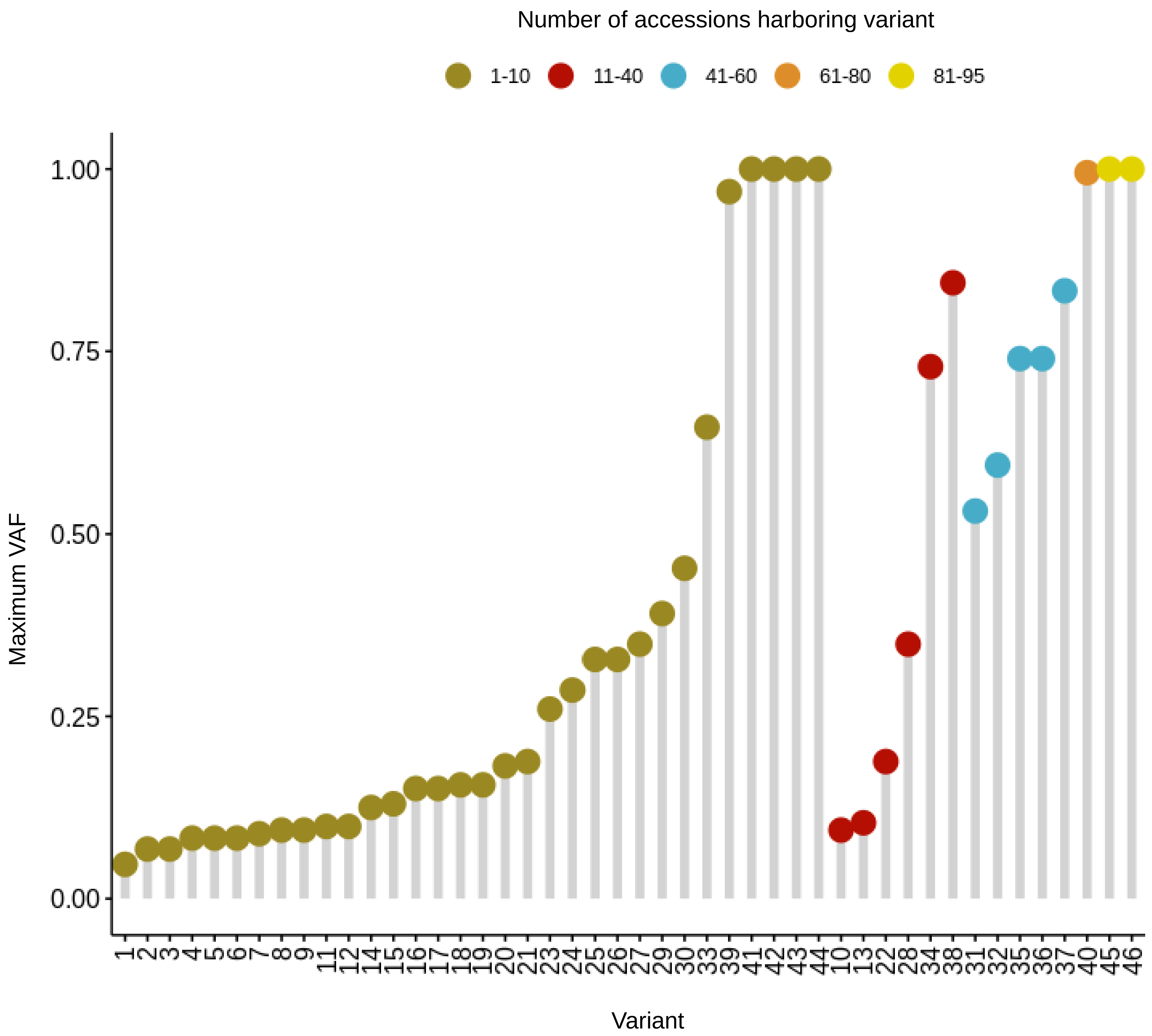

### Supplemental Figure 5

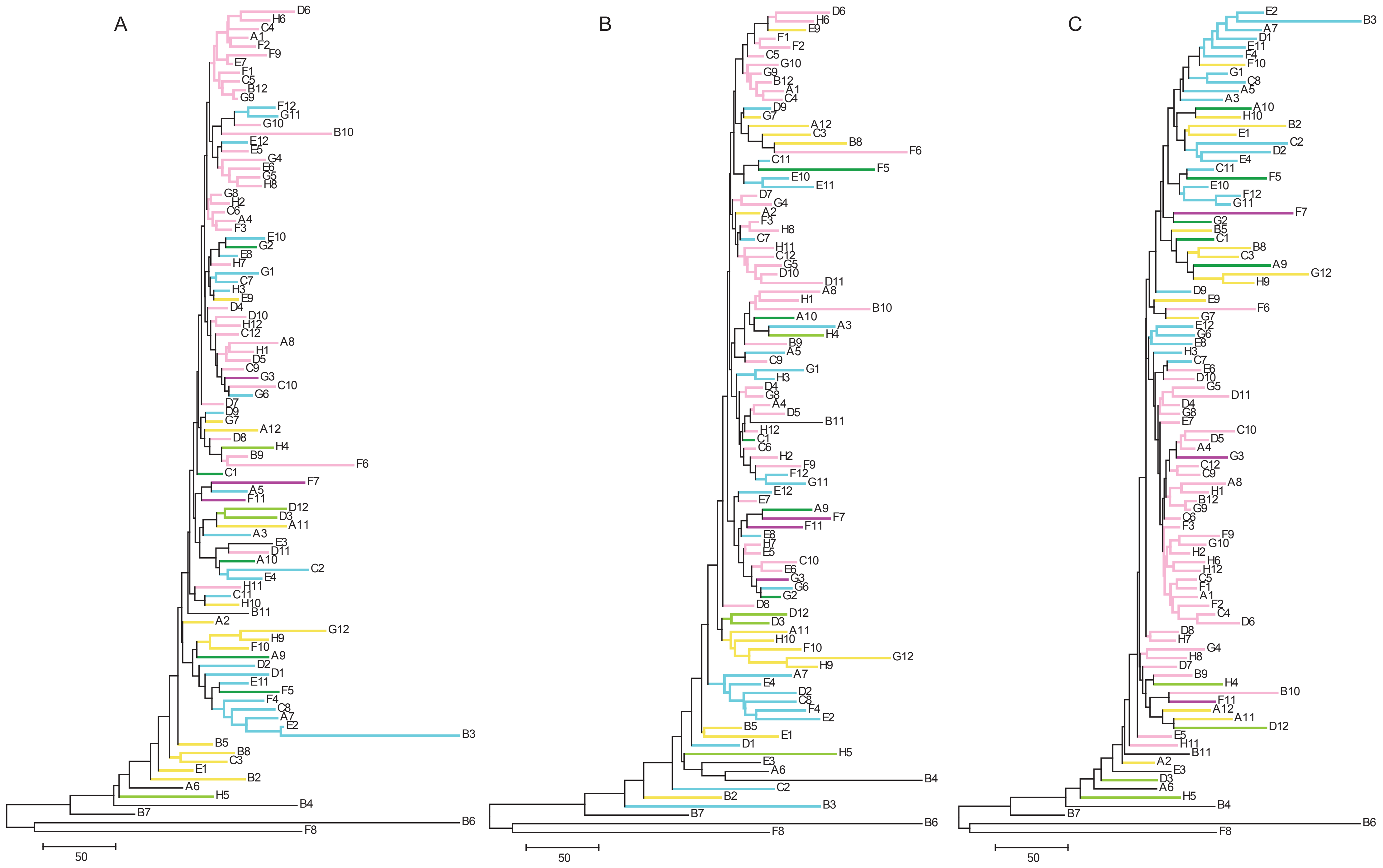

### Supplemental Figure 6

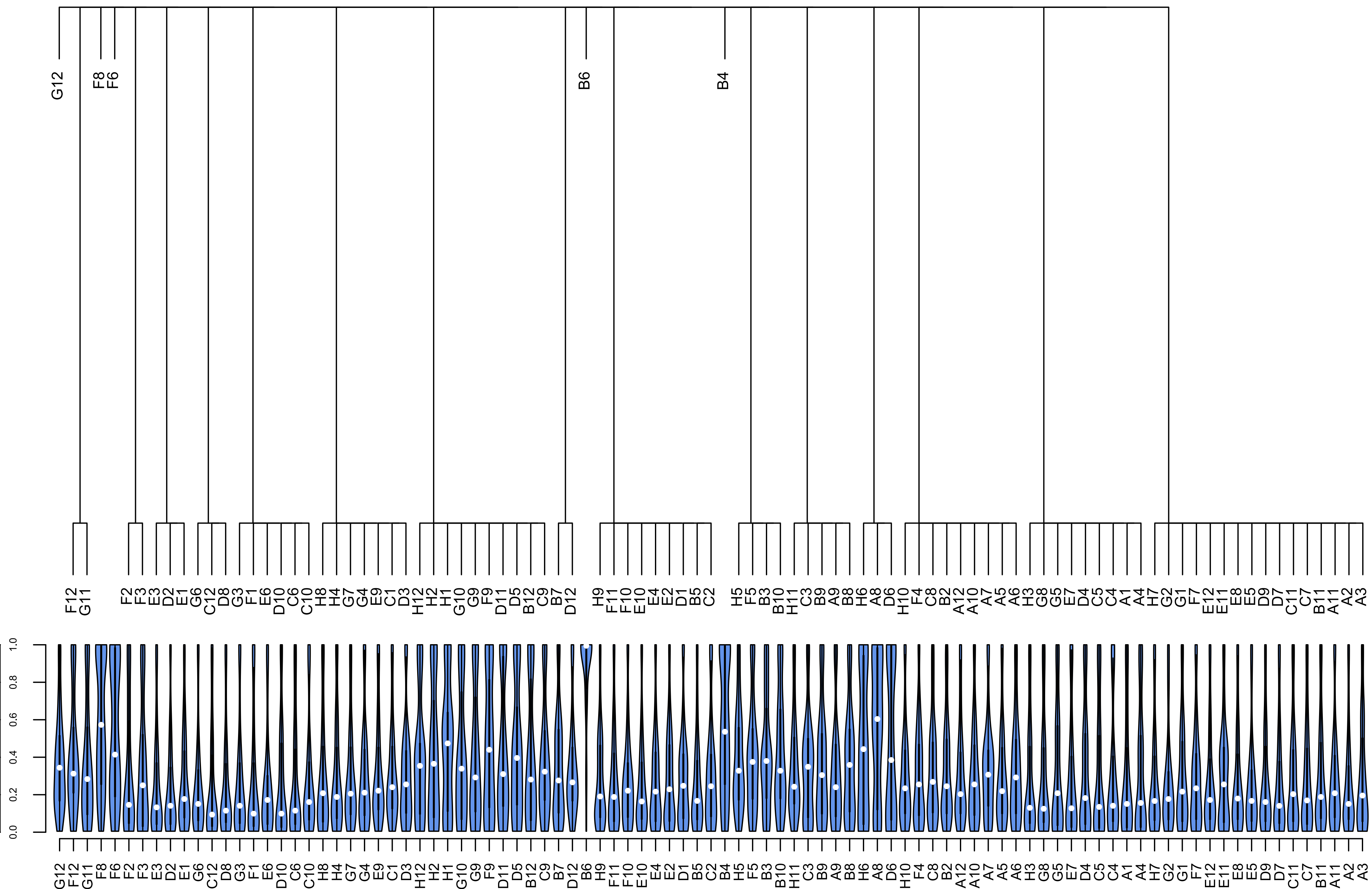

### Supplemental Figure 7

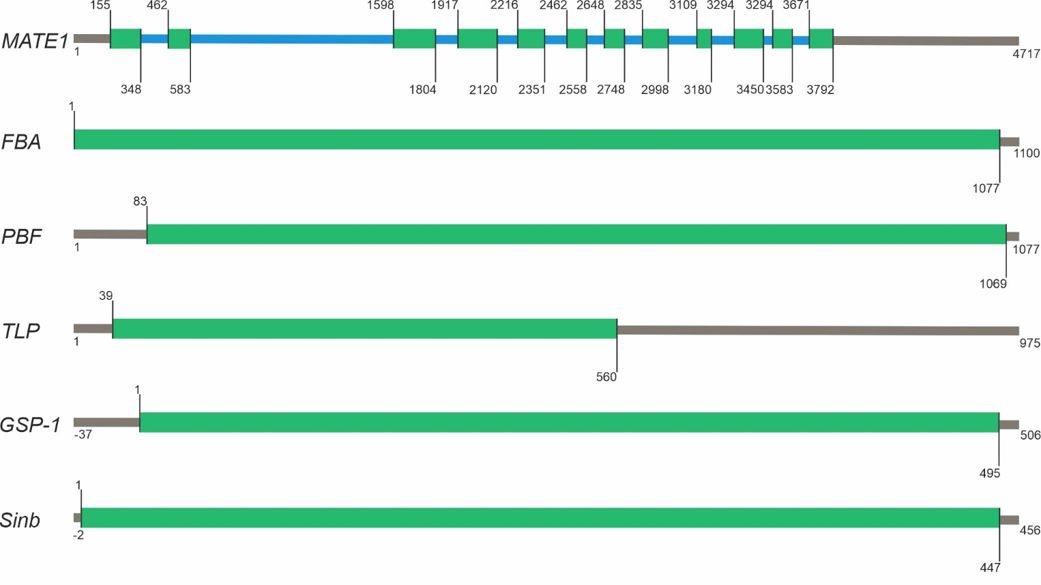
